# Different organic manure sources and NPK fertilizer on soil chemical properties, growth, yield and quality of okra

**DOI:** 10.1101/2020.06.01.127498

**Authors:** Aruna Olasekan Adekiya, Wutem Sunny Ejue, Adeniyi Olayanju, Oluwagbenga Dunsin, Christopher Muyiwa Aboyeji, Charity Aremu, Kehinde Adegbite, Olanike Akinpelu

## Abstract

Use of organic manures to meet the nutrient requirement of crop would be an inevitable practice in the years to come for sustainable agriculture since organic manures generally improve the soil physical, chemical and biological properties. Hence, field experiments were carried out in 2017 and 2018 to compare the impact of different organic manures and NPK fertilizer on soil properties, growth, yield, proximate and mineral contents of okra (*Abelmoschus esculentus* L.). The treatments each year 2017 and 2018 consisted of: rabbit manure, cow dung, poultry manure, green manure [Mexican sunflower (*Tithonia diversifolia* Asteraceae)], pig manure, NPK 15-15-15 fertilizer applied at 120 kg N ha^−1^ and a control (no manure/ inorganic fertilizer). The seven treatments were laid out in a randomized complete block design with three replication. Organic manures and NPK fertilizer increased the soil organic matter (OM), N, P, K, Ca and Mg (NPK fertilizer did not increase OM, Ca and Mg significantly), growth, yield, minerals, protein, ash, carbohydrate and mucilage contents of okra fruit as compared with control. Organic manures improved okra yield compared with NPK fertilizer. Okra growth and yield parameters were significantly higher in 2018 compared with 2017. Control, rabbit manure, cow dung, poultry manure, green manure, pig manure and NPK fertilizer in 2018 increased the pod yield of okra by 9.7%, 35.3%, 57.9%, 36.2%, 39.2%, 45.5% and 3.2%, respectively compare with the same treatment in 2017. Amongst various organic manures, poultry manure produced significantly higher plant growth, yield, mineral and proximate composition of okra because of its high soil chemical properties which could be related to its lowest C: N ratio, lignin and lignin: N ratio. Results also showed that okra grown during high intensity rainfall has higher yield but with reduced quality except its mucilage content. Therefore, planting of okra with poultry manure under moderate rainfall will enhance the health benefit from the fruit, however, those that desire its mucilage content planting during high rainfall is recommended.

## Introduction

Okra (*Abelmoschus esculentus* L.) is an important tropical and subtropical vegetable crop grown for its fresh leaves, buds, flowers, pods, stems and seeds. The fresh pods can be eaten as vegetables in form of salads, in soups and stews or boiled. Okra fruits contain mucilage upon cooking. The mucilage has medical importance as it is used as a plasma replacement or blood volume expander and binds cholesterol and bile acid carrying toxins dumped into it by the liver ^[1]^. Okra fruits contain fiber, vitamin C, folate and antioxidants ^[2]^. The seeds contain oil that is edible by man as well as in soap industry ^[3]^.

Despite the enormous potentials of okra fruits production, its yield per hectare and quality in Nigeria had been greatly hampered by the low fertility status and organic matter contents of the soils which translate into low productivity and consequently reducing income for the farmer ^[4]^. According to Adekiya et al ^[4]^. the yield of okra in Nigeria is currently very low about 2.7 t ha^−1^ owing to low native soil fertility status among other factors. Lack of sufficient amounts of nutrients result in poor performance of the crop with growth been affected resulting to low yield. It has been reported that the maintenance of soil organic matter (OM) is the basis of sustainable crop production in Nigeria and other tropical countries ^[5]^. Hence, there is need to improve the fertility of the soil for continuous and increased crop production.

In years to come, utilization of organic manure to meet crop nutrient requirement will be an unavoidable practice to enhance sustainable agriculture, this is because, the physical, chemical and biological properties of soil is generally improved by the addition of organic manures which in turn enhances crop productivity and maintains the quality of crop produce ^[6]^. Although, in comparison to inorganic fertilizers, organic manures contain smaller quantities of plant nutrients. The use of inorganic fertilizer to increase yield has been found to be effective as a short-term solution but demands consistent use on a long-term basis. The high cost of inorganic fertilizers makes it uneconomical and out of reach to poor farmers and it is also undesirable due to its hazardous environmental effects ^[7]^. Therefore, it is essential to investigate the use of locally sourced organic materials which are environment friendly, cheap and probably an effective way of improving and sustaining the productivity of soils and arable crops such as okra.

Okra yield responses to organic and inorganic fertilizers have been reported by several workers ^[4], [8], [2], [9], [10]^. However, since organic and in organic fertilizers contain different chemical composition and quality they therefore may react differently when applied to the soil with respect to soil chemical properties, crop yield and quality. This aspect needs investigation especially in Nigeria where such data on the effects of different organic and inorganic amendments on the mineral and proximate contents are lacking. Available work dealt with organic amendment on the yield of okra without putting into consideration the quality of okra fruits. In Sokoto northern Nigeria ^[10]^, investigated different sources of organic manure (cow, sheep and poultry manure) on growth and yield of okra. Their results revealed that poultry manure promotes higher growth and yield of okra compared with cow and sheep manure.

Fagwalawa and Yahaya ^[11]^ investigated the effects of Sheep, cow and poultry manures and their combinations on the growth and yield of okra, their result also revealed that poultry manure has the highest yield. Also in Malaysia ^[9]^, six different treatments (no fertilizer, NPK fertilizer, poultry manure, rat manure, goat manure and rabbit manure) were investigated on the growth and yield of okra, According to the study, application of poultry manure significantly increased the growth and yields performances on okra compared to other types of organic fertilizers. To have a holistic approach on the response of organic manures on soil chemical properties and okra performance, data on the quality of such okra fruit is imperative. Therefore, the study is to compare the impact of different organic manures and NPK fertilizer on soil properties, growth, yield, proximate and mineral contents of okra grown in derived savanna zone of Nigeria.

## Material and methods

### Site description, treatments and experimental layout

During 2017 and 2018 cropping seasons, okra was grown in a field experiment at the Landmark University Teaching and Research Farm, Omu-Aran, Kwara state (8°9’N, 5°61’E), with altitude of 562 m above sea level. The rainfall pattern was bimodal with peaks in August (Table 1). The total annual rainfall in the area was about 1238 mm in 2017 with mean air temperature of 28.7°C and mean relative humidity of 83.9%. In 2018, the total annual rainfall in the area was about 1428 mm with mean air temperature of 30.0°C and mean relative humidity of 77.4%. The soil at the site of the experiment is an Alfisol classified as Oxic Hap-lustalf or Luvisol^[2]^. The experimental site falls under the derived savanna agro-ecological zone of North-Central Nigeria. Weeds in the experimental soil before cultivation included Mexican sunflower (*Tithonia diversifolia* Asteraceae) and Guinea grass (*Panicum maximum* Jacq). Different sites were used for the experiment in 2017 and 2018, but the two sites were very close to each other.

**Table 1:** Meteorological data of the study area.

| 2017 | Jan. | Feb. | Mar. | Apr. | May | Jun | Jul | Aug | Sep | Oct | Nov | Dec |
| --- | --- | --- | --- | --- | --- | --- | --- | --- | --- | --- | --- | --- |
| Rainfall (mm) | 0 | 0 | 97.28 | 92.46 | 158.24 | 174.49 | 71.11 | 179.33 | 231.39 | 119.88 | 0 | 114.04 |
| Relative humidity (%) | 54.1 | 44.9 | 69.7 | 81.0 | 85.3 | 89.9 | 91.9 | 93.4 | 92.3 | 85.2 | 71.3 | 56.9 |
| Mean Temp (°C) | 29.1 | 31.2 | 31.9 | 29.9 | 28.9 | 27.9 | 26.9 | 26.3 | 26.5 | 27.8 | 29.9 | 28.4 |
| 2018 | Jan. | Feb. | Mar. | Apr. | May | Jun | Jul | Aug | Sep | Oct | Nov | Dec |
| Rainfall (mm) | 0 | 12.19 | 157.73 | 96.26 | 214.88 | 228.6 | 160.02 | 95.25 | 248.66 | 195.07 | 19.55 | 0 |
| Relative humidity (%) | 34.3 | 63.7 | 78.8 | 85.2 | 87.5 | 90.2 | 92.7 | 92.7 | 91.3 | 88.4 | 76.4 | 47.1 |
| Mean Temp (°C) | 26.6 | 29.6 | 29.3 | 29.2 | 28.1 | 27.7 | 26.4 | 26.2 | 26.4 | 27.5 | 28.8 | 28.9 |

Source: Meteorological unit, Teaching and Research Farm, Landmark University, Omu-Aran, Kwara State, Nigeria.

The treatments each year 2017 and 2018 consisted of: rabbit manure, cow dung, poultry manure, green manure [Mexican sunflower (*Tithonia diversifolia* Asteraceae)], pig manure, NPK 15-15-15 fertilizer and a control (no manure/ inorganic fertilizer). Both the organic manures and NPK fertilizer were applied at the rate of 120 kg N ha^−1 [12]^. This was equivalent to 800 kg ha^−1^ for NPK 15-15-15, 11.9 t ha^−1^ for rabbit manure, 5.5 t ha^−1^ for cow dung, 4.1 t ha^−1^ for poultry, 4.8 t ha^−1^ for green manure and 5.6 t ha^−1^ for pig manure. The seven treatments were laid out in a randomized complete block design (RCBD) with three replications. Each block consisted of 7 plots measuring 3 × 2 m with 1m block and 0.5m between plots.

### Land preparation, incorporation of manures, sowing of okra seeds and application of NPK fertilizer

The experimental field was cleared manually using cutlass and thrashed removed from the site before mechanical (ploughing and harrowing) land preparation after which plots were marked out to the require plot size of 3 × 2 m. A hand-held hoe was used to incorporate the manures into the soil to a depth of approximately 20 cm. All animal manures were obtained from the livestock section of Landmark University Teaching and Research Farm. Fresh top Mexican sunflower was collected from a nearby farm and hedge containing green tender stems and the leaves. The plots were allowed for 4 week before sowing of okra seeds. Sowing of okra variety NHAe-47-4 was done late April for both 2017 and 2018 croppings. Two seeds of okra were sown per hole at inter-row spacing of 0.6 m and 0.6 m intra-row spacing manually giving a plant population of 27,778 plants ha^−1^. At two weeks after sowing, thinning to one plant per stand was done and this was followed by manual weeding using hand hoe before treatment application. Subsequent weeding was done as needed. NPK 15-15-15 fertilizer was applied by side placement at about 8-10 cm away from the sown seeds two weeks after sowing. Insect pests were controlled by spraying cypermethrin weekly at the rate of 30 ml per 10 L of water from 2 weeks after sowing till 4 weeks after sowing.

### Determination of soil properties

Before the start of the experiment, soil samples from topsoil (0 – 15 cm) were taken from random spots in the study area and were bulked together, air-dried and sieved using a 2-mm sieve and their physical and chemical characteristics were determined^[2]^. Textural class of the soils were determined by the method of ^[13]^. Soil organic carbon (OC) was determined by the procedure of Walkley and Black using the dichromate wet oxidation method^[14]^. Total N was determined by the micro-Kjeldahl digestion method^[15]^. Available P was determined by Bray-1 extraction followed by molybdenum blue colorimetry^[16]^. Exchangeable K, Ca, and Mg were extracted using 1M ammonium acetate^[17]^. Thereafter, concentration of K was determined on a flame photometer, and Ca and Mg were determined by EDTA titration method. Soil pH was determined using a soil-water medium at a ratio of 1:2 with a digital electronic pH meter. At the termination of the experiment in 2017 and 2018, soil samples were also collected (on plot basis) and similarly analysed for soil chemical properties as described above. Also at the end of the experiment each year, soil bulk density were determined all plots. Five undisturbed samples (0.04 m diameter, 0.15 m high) were collected at 0 – 0.15 m depth from the centre of each plot at random and 0.15 m away from each okra plant using core steel sampler^[4]^. The samples were used to evaluate bulk density after oven-drying at 100^°^C for 24 h^[4]^

### Determination of growth and yield parameters

Collection of data for growth (plant height and leaf area) was done at mid-flowering of okra plant (about 47 days after sowing). Plant height was determined by the use of meter rule while leaf area was determined by using the model {LA=0.34(LW)^1.12^} developed by^[18]^, where LA = leaf area, L= leaf length and W = leaf width. Edible pods were harvested at four day intervals, counted and weighed. Pod weight was evaluated based on the cumulative harvests per plot. For 2017 cropping of okra, fruits were harvested until November 2017 while for 2018 okra cropping, fruits were harvested until July 2018.

### Chemical analysis of okra fruits and organic manures

At harvest in 2017 and 2018, eight okra fruits of uniform sizes were randomly collected from each plot and analyzed for mineral contents according to methods recommended by the Association of Official Analytical Chemists^[19]^. One gram of each sample was digested using 12 cm^−3^ of the mix of HNO_3_, H_2_SO_4_, and HCLO_4_ (7:2:1 v/v/v). Contents of N, P, K, Ca and Mg were determined by atomic absorption spectrophotometry. Lignin content of the organic amendments was determined from acid-free detergent fiber using the method described by ^[20]^.

Samples of okra fruits from each plot was taken for proximate analysis. The ash, crude protein, crude fat and carbohydrate contents of the okra fruits were determined using standard chemical methods described by Association of Analytical chemists ^[19]^. Samples of okra fruits were sliced with a knife and blended. After blending, it was diluted with ten times its weight with water (1:10). The viscous solution was separated from the debris using fine cloth ^[21]^. Mucilage of the extracted viscous liquid was measured using viscometer.

About 2 g of each organic manures used was collected and analysed for N, P, K, Ca, and Mg as described by ^[22]^. N was determined by the micro-Kjeldahl digestion method. Samples were digested with nitricperchloric-sulphuric acid mixture for the determination of P, K, Ca, and Mg. Phosphorus was determined colorimetrically using the vanadomolybdate method, K was determined using a flame photometer and Ca and Mg were determined by the EDTA titration method ^[23]^.

### Statistical analysis

The data collected were subjected to statistical analysis of variance (ANOVA) using the Genstat statistical package ^[24]^ and treatment means were separated using Duncan Multiple Range Test (DMRT) at 5% probability level.

## Results

### Pre-planting chemical and physical analysis of the experimental field and chemical composition of organic manures used for the experiment

The result of the pre-planting chemical and physical analysis of the experimental field (Table 2), indicated in both years that the soil was sandy loam and slightly acidic with a pH of 6.10 and 6.20 respectively in 2017 and 2018. The soil organic matter, total N available P and exchangeable Ca and K were low with both a little below the critical levels of 3.0 %, 0.20 %, 10.0 mg kg^−1^, 2.0 cmol kg^−1^and 0.15 cmol kg^−1^ respectively in both cropping seasons of 2017 and 2018 ^[25]^. Exchangeable Mg was adequate in 2017 but low in 2018. Analysis of the soil amendments used for this experiment is shown in Table 3 with poultry manure having the highest percentage of N, P, K, Ca, Mg and the lowest lignin, lignin: N ratio, C: N ratio and organic C.

**Table 2:** Pre-planting physico-chemical characteristics of the soil (0 – 15 cm) at the experimental sites in 2017 and 2018.

| Soil properties | 2017 | 2018 |
| --- | --- | --- |
| pH (H<sub>2</sub>O) | 6.10 | 6.20 |
| OM (%) | 2.56 | 2.44 |
| Total N (%) | 0.10 | 0.15 |
| Available P (mg/kg) | 9.50 | 8.90 |
| Exchangeable cations (cmol kg<sup>-1</sup>) |  |  |
| K | 0.15 | 0.14 |
| Ca | 1.80 | 1.90 |
| Mg | 0.41 | 0.38 |
| Particle size distribution (%) |  |  |
| Sand | 76 | 74 |
| Silt | 13 | 15 |
| Clay | 11 | 11 |
| Textural class | Sandy loam | Sandy loam |

**Table 3:**
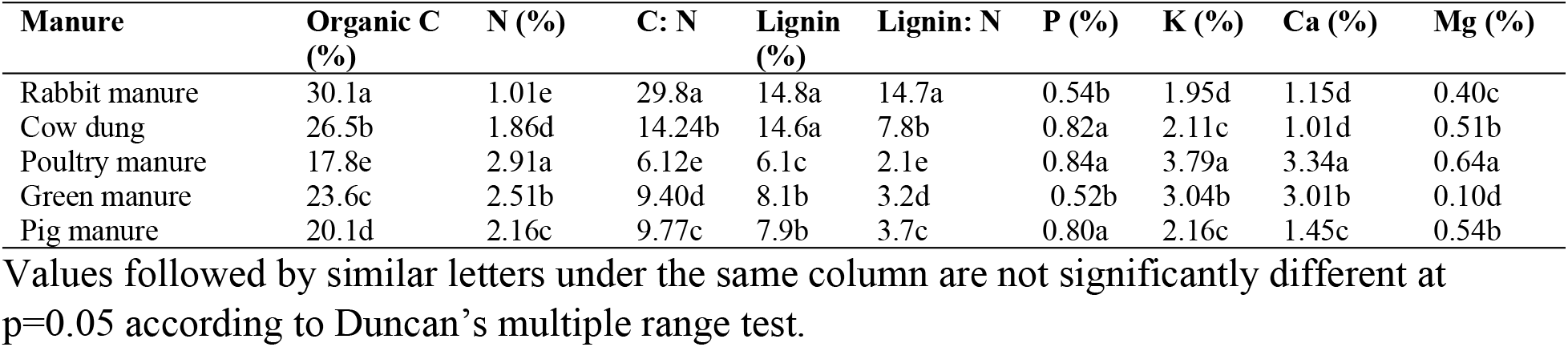
Analysis of organic manures.

### Response of soil chemical properties and bulk density to different organic manure sources and NPK fertilizer

Table 4 shows the result of the effect of different organic amendments and NPK fertilizer on soil chemical properties. In both years, all organic sources of soil amendment increased soil OM, N, P, K, Ca and Mg significantly (p<0.05) with respect to the control. NPK fertilizer did not increase OM, Ca and Mg contents of the soil relative to the control but increased N, P and K. Among organic manure sources, poultry manure has the highest values of all soil nutrients (except SOM) which was closely followed by green manure treatment with rabbit manure having the least values. The order of SOM among organic manures was: Rabbit manure > cow dung > Pig manure > green manure > Poultry manure. All organic manure sources increased soil N, P and K compared with NPK fertilizer. In both years, the control has the highest value of pH while NPK fertilizer has the least value, there were no significant differences between the pH values of control, rabbit manure, cow dung, poultry manure, green manure and pig manure. Also there were no significant differences in the pH values between organic manures and NPK fertilizer, but there was a significant differencs between the pH values of the control and NPK fertilizer. Soil chemical properties in year 2018 were increased compared with year 2017 except soil pH. Soil bulk density in both years was significantly reduced in organic manure soils compared with the control and NPK fertilizer which has similar values. All organic manures have statistically similar values of bulk density. The interactions between year and amendment (Y × M) were significant for all soil properties except soil pH and bulk density

**Table 4:**
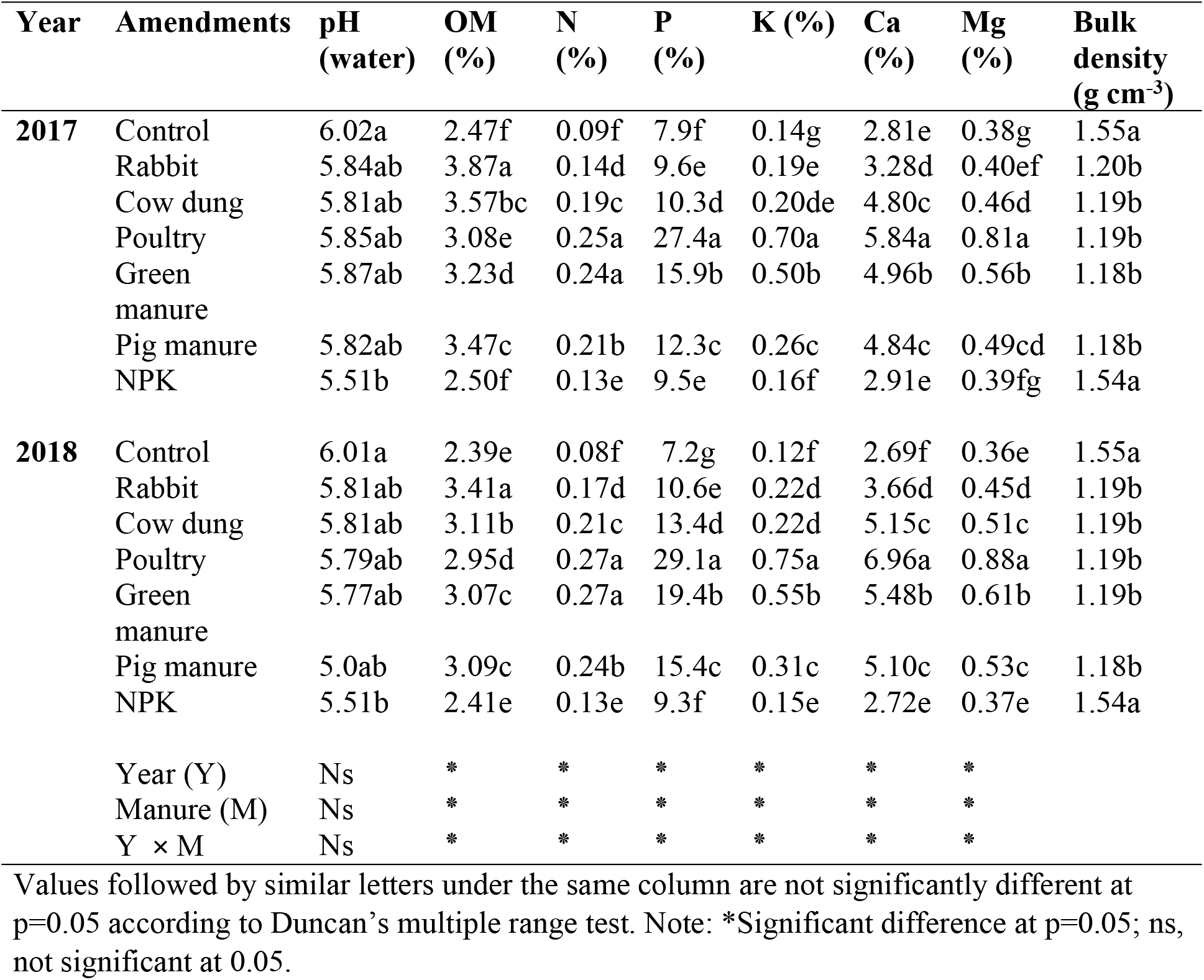
Post-Harvest soil chemical properties.

### Response of okra growth and yield to different organic manure sources and NPK fertilizer

The results of the response of okra growth and yield to different organic manure sources and NPK fertilizer are presented in Table 5. In 2017 and 2018, all amendments (both organic and inorganic) increased the growth (plant height and leaf area) and yield (okra pod weight and number of pods per plant) of okra compared with the control. Also in both years, poultry manure has the highest values followed by green manure. The order of okra pod yield was: poultry manure > green manure > pig manure > cow dung > NPK fertilizer = rabbit manure > control. Okra growth and yield parameters were significantly higher in 2018 compared with 2017. Control, rabbit manure, cow dung, poultry manure, green manure, pig manure and NPK fertilizer in 2018 increased the pod yield of okra by 9.7%, 35.3%, 57.9%, 36.2%, 39.2%, 45.5% and 3.2%, respectively compare with the same treatment in 2018. The interactions between year and amendment (Y × M) were significant for all growth and yield parameters.

**Table 5:** Growth and yield parameters.

| Year | Amendments | Yield (t ha<sup>-1</sup>) | No. of pods/<br>plant | Plant height<br>(m) | Leaf area<br>(m<sup>2</sup>) |
| --- | --- | --- | --- | --- | --- |
| **2017** | Control | 3.1f | 10g | 0.29f | 0.46e |
|  | Rabbit | 3.4e | 12f | 0.37e | 0.54d |
|  | Cow dung | 3.8d | 14de | 0.41d | 0.57d |
|  | Poultry | 5.8a | 23a | 0.56a | 0.81a |
|  | Green manure | 5.1b | 21b | 0.51b | 0.71b |
|  | Pig manure | 4.4c | 17c | 0.47c | 0.64c |
|  | NPK | 3.5e | 13ef | 0.38e | 0.55d |
| **2018** | Control | 3.4g | 12h | 0.30f | 0.50f |
|  | Rabbit | 4.6e | 15e | 0.41de | 0.60e |
|  | Cow dung | 5.2d | 16de | 0.44d | 0.65de |
|  | Poultry | 7.9a | 26a | 0.62a | 0.94a |
|  | Green manure | 7.1b | 24b | 0.55b | 0.80b |
|  | Pig manure | 6.4c | 18c | 0.50c | 0.71c |
|  | NPK | 3.6f | 14fg | 0.39e | 0.60e |
|  | Year (Y) | * | * | * | * |
|  | Manure (M) | * | * | * | * |
|  | Y × M | * | * | * | * |

Values followed by similar letters under the same column are not significantly different at p=0.05 according to Duncan's multiple range test. Note: \*Significant difference at p=0.05

### Response of okra quality to different organic manure sources and NPK fertilizer

The effect of different organic manure sources on proximate and mineral content of okra in 2017 and 2018 are shown in Table 6. In 2017 and 2018 different organic manure sources and NPK fertilizer increased protein, ash, carbohydrates, mucilage N, P, K Ca and Mg contents of okra fruits compared with the control. Fat contents of okra fruits were reduced with different organic sources and NPK fertilizer compared with the control. Among different amendment applied, poultry manure had the highest values of protein, carbohydrate, mucilage, N, P, K, Ca and Mg contents. Green manure had statistically similar value with poultry manure in protein, ash, fat, carbohydrate, mucilage, N, P and Mg contents in 2017 and in 2018, ash, fat, carbohydrate, mucilage, N and P were statistically similar. Poultry manure and green manure increased protein, carbohydrate, mucilage, N, P, K, Ca, Mg and reduced fat compared with NPK fertilizer. 2018 cropping of okra reduced protein, ash, fat, carbohydrate, N, P, K, Ca, Mg and increased mucilage contents in the fruits compared with 2017. The interaction (Y × M) was also significant for proximate and mineral contents of the okra fruit.

**Table 6:**
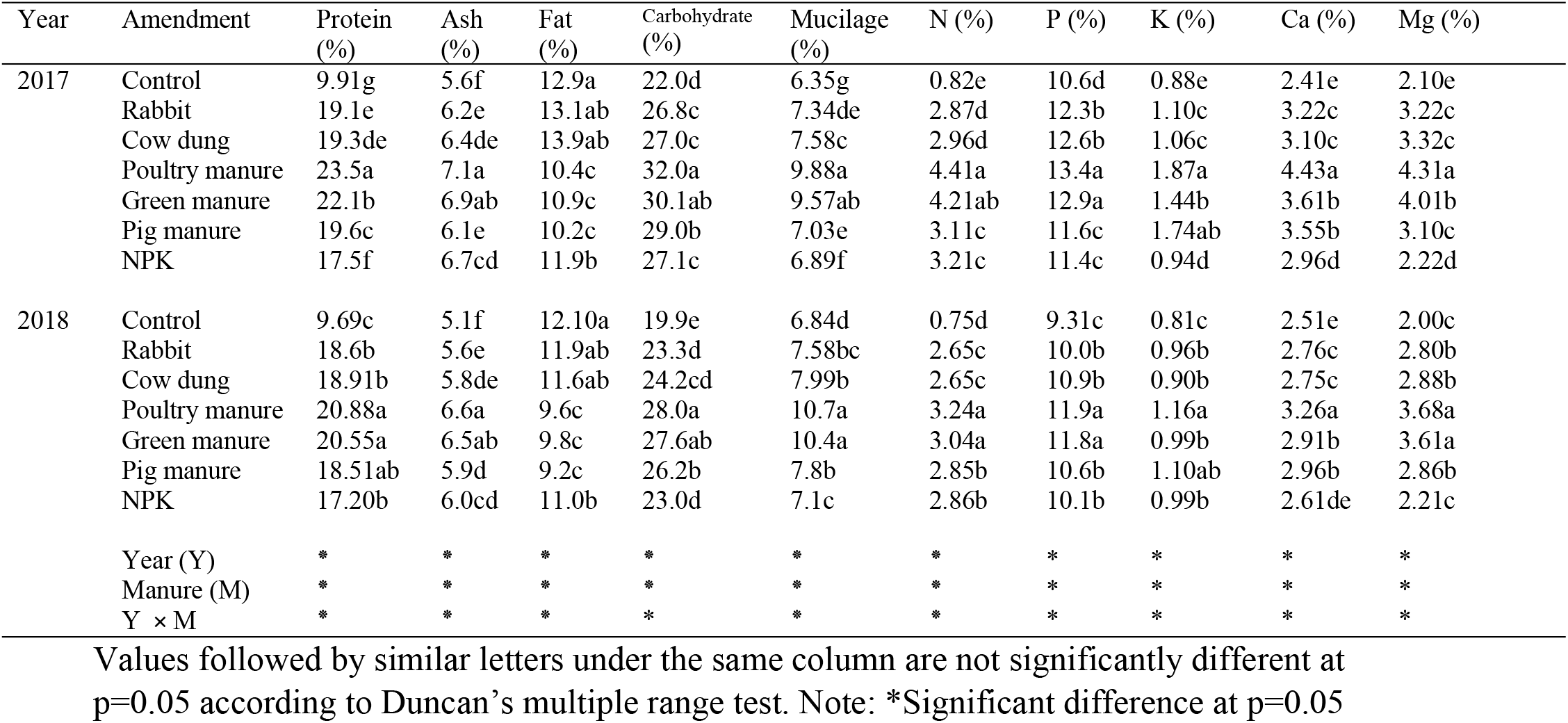
Proximate and mineral contents of okra.

## Discussion

The different organic manures increased the soil organic matter (SOM), N, P, K, Ca and Mg contents of the soil compared with control. This result is consistent with the analysis recorded for the amendments in the present study that they contain these nutrients and also attested to the fact that the soil was deficient in these nutrients (Table 2). These nutrients were released into the soil as the organic manures were decomposed. Studies by ^[26] & [27]^ have shown that animal manures and green manure increased soil OM, N, P and CEC and this was attributed to the availability and adequate supply of organic matter. The slightly lower pH of organic amended soil compared with the control could be due to the fact that during microbial decomposition of the incorporated manures, organic acids may have been released, which neutralized the alkalinity of the manures, thereby lowering the pH of the soil below their initial values. Adekiya et al.^[4]^ observed a similar trend in their work on organic amendments of soils. NPK fertilizer has the lowest pH as a result of leaching of bases from soil surface. Poultry manure significantly produced the highest soil chemical properties, this could be related to its lowest C:N ratio, lignin and lignin: N ratio (Table 3) which favours quick mineralization and release of nutrients to the soil compared with other soil amendments. Consequently, the quality of an organic amendment is defined in term of the relative content of nutrients (especially nitrogen), lignin, lignin: N and the C: N ratio ^[27]^. Rabbit manure increased SOM compared with other organic manures. The increase in SOM of rabbit manure could be related to its high C: N and lignin content. Plant constituents, such as lignin retard decomposition. Organic materials with high C: N and lignin generally would favor nutrient immobilization, organic matter accumulation and humus formation ^[27]^.

In both years, poultry manure increased N, P, K, Ca and Mg compared with other manures. This was due to the fact that poultry manure was low in C: N ratio, lignin and lignin/N and high in N, P, K, Ca and Mg compared with other manures. Due to high quality of this amendment, it decomposes quickly and release nutrients to the soil. The significant increase in soil nutrients especially OM, N, P and K in organic amendments compared with NPK fertilizer was due to leaching in NPK fertilizer treated plots. The reduction in Ca and Mg in NPK fertilizer plots compared with organically amended plot was due to the fact that NPK fertilizer did not contain Ca and Mg. The reduction of soil bulk density observed in both years with organic manures compared with control and NPK fertilizer could be attributed to increase in soil organic matter resulted from the degraded organic residues by soil microorganisms. Organic matter is known to improve soil structure, aeration and reduce soil bulk density ^[28]^.

The decreasing order of okra yield were poultry manure > green manure > pig manure > cow dung > NPK fertilizer = rabbit manure > control. The positive effect of organic manure on growth and okra yield could be due to the contribution made by amendments to fertility status of the soils as the soils were low in organic carbon content. Manure when decomposed increases both macro and micro nutrients as well as enhances the physio-chemical properties of the soil for the betterment of okra growth.

Okra grown on poultry manure performed better in terms of growth and yield compared with other sources of organic soil amendment and NPK fertilizer. This could also be related to low C:N ratio, lignin and lignin/N values These attributes of poultry manure will lead to fast mineralization and early release of nutrients to a short gestation crop like okra, hence there was a boost in the morphological growth of the plant which translate to greater yield compared with other amendments. Wolf and Snyder ^[29]^ reported that C: N ratio of organic materials markedly influences the decomposition rate and the mineralization of N because N determines the growth and turnover of the microorganisms that mineralize organic carbon.

The reduced growth and yield of okra in plots treated with other manures in comparison with poultry manure could be as a result of immobilization of soil nutrients which occurs when soils are treated with animal manure of high lignin contents which results from the feed eaten by the animal ^[30]^.

Higher yields were observed in poultry manure plots compared with inorganic fertilizer because, the nitrogen content of poultry manure is released to the soil gradually and steadily over longer time for the growth of the plant compared with nitrogen from NPK fertilizer which is prone to losses by run-off, volatilization, leaching and/or denitrification. Poultry manure has been said to be a better soil amendment compared with chemical fertilizers because of the greater capability of poultry manure to preserve its N^31^. The superior N supply by poultry manure during okra cropping in this experiment may be the reason for better growth and yield of okra in plots with poultry manure. The obtained results corroborated the finding of^32^ that poultry manure increased the height of okra relative to other amendments.

The better performance of okra under NPK fertilizer plots compared with the control was due to release of nutrients (N, P and K) from the fertilizer which are absorbed by the okra plants. Okra growth and yield in second year (2018) was better than that of first year (2017). This was due to the differences in the amount of rainfall between the two years (Table 1). Year 2017 had 1238 mm of rainfall while it was 1428 mm in 2018. There was higher moisture in the first few weeks (month of May) of incorporation of the manures in 2018 compared with 2017 which may have led to better and quicker decomposition of the organic materials in 2018. Soil biological activities which causes degradation of organic materials is severely limited during limited moisture, but with the onset of the rains (2018), there is a flush in microbial activity ^[33]^.

The fact that organic manures and NPK fertilizer increased okra mineral contents compared with the control was attributed to increased availability of the nutrients in soil as a result of the mineralization of the manures leading to increased uptake by okra plants. In both years, the correlation coefficient between soil and okra fruit N, soil and okra fruit P, soil and okra fruit K, and okra fruit Ca, soil and okra fruit Mg were all significant with R values of 0.83, 0.71, 0.66 0.81 and 0.88, respectively at *P* < 0.05. Poultry manure had the highest values of N, P, K, Ca and Mg in the okra fruit compared with other amendment and NPK fertilizer. This result is also consistent with the soil chemical properties and growth and yield for this treatment. The poultry manure may have improved the availability of nutrients to the crop by enhancing the mineralization and supply of readily available nutrients to the soil ^[34]^.

The increased nitrogen to the soil by the incorporation of organic manures increased the nitrogen uptake by the okra fruit thereby increasing protein. This explains the reasons for increase in crude protein values between organic amendment and control and NPK fertilizer treatment. Poultry manure has the highest value of crude protein, this is also consistent with its high soil N level (Table 4). Nitrogen is a major constituent of chlorophyll, protein, amino acids, various enzymes, nucleic acids and many other compounds in the cell of plants ^[35]^. The crude protein contents in all the organically amended soils were above the critical 13-17% ^[36], [37]^. Therefore the organic materials sustained good nutritive quality of okra as opposed to NPK and control. The ash content in poultry manure treatment was significantly higher possibly because of the balanced nutrient in the manure, unlike NPK with inferior contents of N, P, and K and the control with lower concentration of nutrients. In this study, poultry manure gave the highest N, P and K concentrations of 3.04%, 11.9% and 1.16% in 2017 and 4.41% 13.4% and 1.87% in 2018 (Table 6) respectively. The fat content of poultry manure was lowered compared with others, this was due to its high protein content of poultry manure. It has also been reported ^[38]^ that there is a negative correlation between fat and protein content. High nitrogen application reduced fat and increased protein content. Organic amendments and NPK fertilizer increased the mucilage contents of okra compared with the control. This could be as a result of increase in NPK from these amendments. This could be attributed to the increase in D- galactose, L. rhamnose and D- galacturonic acid contents in okra fruits by the application of nutrients through organic and inorganic sources which might have resulted in increase of mucilage content ^[39]^. The mucilaginous polysaccharide in the okra is rich in uronic acid (65%) and consists of rhamnose, galactose, glucose, galacturonic acid and glucuronic acid in addition to 3.7% acetyl groups ^[40]^. Ahmad et al.^[40]^ also reported that compost and NPK fertilizer application increased the mucilage of borage plant. The highest value of mucilage in poultry manure could be related to increased soil nutrients compared with other treatments. Okra produced under organic amendments has better qualities (N, P, K, Ca, Mg, protein, ash and mucilage) compared with NPK fertilizer and control. This is because organic manures not only increase soil nutrients but also improves the physical by prevention of erosion and leaching of nutrients (in this experiment reduce bulk density) and biological properties ^[41]^. Organic manures also contains both micro and macro nutrients unlike NPK fertilizer that contains only N, P and K. Lumpkin ^[42]^ also, was of the opinion that organically produced vegetables were of higher qualities than those produced using conventional methods. Cropping of okra in 2018 reduced protein, ash, fat, carbohydrate, N, P, K, Ca and Mg contents of okra fruit compared with 2017 cropping. This could be adduced to high rainfall in 2018 compared with 2017. High rainfall has been reported to reduce the nutrient and proximate content of vegetables ^[43]^. Climatic conditions have a great effect on the concentration of mineral in plants. Variation in temperature and rainfall have been reported to influence the chemical composition in plants ^[43], [44]^.

## Conclusion

Results of this experiment showed that organic manures and NPK fertilizer increased the soil chemical properties (NPK fertilizer did not increase OM, Ca and Mg significantly), growth, yield, minerals, protein, ash, carbohydrate and mucilage contents of okra fruit as compared with control. Organic manures improved okra yield compared with NPK fertilizer. Amongst various organic manures, poultry manure produced significantly higher plant growth, yield, mineral and proximate composition of okra because of its high soil chemical properties which could be related to its lowest C:N ratio, lignin and lignin: N ratio. Results also showed that okra grown during high intensity rainfall has higher yield but with reduced quality except its mucilage content. Therefore, planting of okra with poultry manure under moderate rainfall will enhance the health benefit from the fruit, however, those that desire its mucilage content planting during high rainfall is recommended.

## Acknowledgements

I would like to thank Landmark University who provided enabling environment in carrying out the experiment and also willing to pay the article processing charges of this paper

## Author Contributions

**Conceptualization:** Aruna Olasekan Adekiya, Oluwagbenga Dunsin, Christopher Muyiwa Aboyeji

**Data curation:** Aruna Olasekan Adekiya, Oluwagbenga Dunsin, Christopher Muyiwa Aboyeji, Wutem Sunny Ejue

**Formal analysis:** Aruna Olasekan Adekiya, Adeniyi Olayanju

**Investigation:** Aruna Olasekan Adekiya, Oluwagbenga Dunsin, Christopher Muyiwa Aboyeji

**Methodology:** Aruna Olasekan Adekiya, Kehinde Adegbite, Charity Aremu, Olanike Akinpelu

**Resources:** Aruna Olasekan Adekiya, Olanike Akinpelu, Wutem Sunny Ejue

**Supervision:** Aruna Olasekan Adekiya, Oluwagbenga Dunsin, Christopher Muyiwa Aboyeji

**Writing – original draft:** Aruna Olasekan Adekiya, Charity Aremu, Adeniyi Olayaju.

